# Combined deletion of free fatty-acid receptors 1 and 4 minimally impacts glucose homeostasis in mice

**DOI:** 10.1101/2020.08.04.236471

**Authors:** Marine L. Croze, Arthur Guillaume, Mélanie Ethier, Grace Fergusson, Caroline Tremblay, Scott A. Campbell, Hasna Maachi, Julien Ghislain, Vincent Poitout

**Author notes:** Corresponding author and reprint requests: Vincent Poitout, DVM, PhD, CRCHUM, 900 rue St Denis, Montréal, QC, H2X 0A9 - CANADA. This work was supported by a Discovery Grant from the Natural Sciences and Engineering Research Council of Canada (RGPIN-2016-03952 to V.P.). M.L.C. was supported by a fellowship from the Société Francophone du Diabète and a postdoctoral fellowship from the Montreal Diabetes Research Center. The authors have no relevant conflict of interest to disclose.

## Abstract

The free fatty-acid receptors FFAR1 (GPR40) and FFAR4 (GPR120) are implicated in the regulation of insulin secretion and insulin sensitivity, respectively. Although GPR120 and GPR40 share similar ligands, few studies have addressed possible interactions between these two receptors in the control of glucose homeostasis. Here we generated mice deficient in *gpr120* (Gpr120KO) or *gpr40* (Gpr40KO), alone or in combination (Gpr120/40KO), and metabolically phenotyped male and female mice fed a normal chow or high-fat diet. We assessed insulin secretion in isolated mouse islets exposed to selective GPR120 and GPR40 agonists singly or in combination. Following normal chow feeding, body weight and energy intake were unaffected by deletion of either receptor, although fat mass increased in Gpr120KO females. Fasting blood glucose levels were mildly increased in Gpr120/40KO mice, and in a sex-dependent manner in Gpr120KO and Gpr40KO animals. Oral glucose tolerance was slightly reduced in male Gpr120/40KO mice and in Gpr120KO females, whereas insulin secretion and insulin sensitivity were unaffected. In hyperglycemic clamps, the glucose infusion rate was lower in male Gpr120/40KO mice but insulin and c-peptide levels were unaffected. No changes in glucose tolerance were observed in either single or double KO animals under high-fat feeding. In isolated islets from wild-type mice, the combination of selective GPR120 and GPR40 agonists additively increased insulin secretion. We conclude that while simultaneous activation of GPR120 and GPR40 enhances insulin secretion ex vivo, combined deletion of these two receptors only minimally affects glucose homeostasis in vivo in mice.

## INTRODUCTION

Insulin resistance and defective insulin secretion are hallmarks of type 2 diabetes (T2D). A number of G protein-coupled receptors (GPCR) that regulate insulin sensitivity and pancreatic β-cell function are validated targets for the treatment of T2D [1]. Among these, the long-chain fatty acid receptors FFAR1 (GPR40) and FFAR4 (GPR120) have been the subject of increasing interest in recent years as their activation has numerous beneficial effects on glucose and energy homeostasis in preclinical T2D models.

In rodents, *gpr120* is expressed in several tissues implicated in energy homeostasis including the central nervous system, enteroendocrine cells, liver, bone, adipose tissue and macrophages [2]. GPR120 activation in adipocytes and macrophages alleviates obesity-induced insulin resistance [3-5] in part via interaction with peroxisome proliferator-activated receptor γ [6]. GPR120 also promotes adipogenesis [7-9] and brown adipose tissue thermogenesis [10,11], regulates food intake [12] and modulates gut endocrine hormone secretion, including ghrelin [13-15] and the incretin hormones glucagon-like peptide-1 (GLP-1) [16], gastric inhibitory polypeptide (GIP) [17], cholecystokinin [18,19] and somatostatin [20]. GPR120 is also expressed in pancreatic islet cells where its activation mitigates β-cell dysfunction [21] and apoptosis [22] and modulates insulin [23-26], glucagon [27], somatostatin [28] and pancreatic polypeptide [29] secretion. Recently, we demonstrated that GPR120 activation potentiates glucose-stimulated insulin secretion (GSIS) and arginine-induced glucagon secretion in part through inhibition of somatostatin release by δ cells [30]. Despite the evidence for a role of GPR120 in the regulation of islet hormone secretion, *gpr120* deficient mice have reportedly normal β-cell function [3,5].

In rodents, *gpr40* is predominantly expressed in enteroendocrine cells and pancreatic β cells [31,32]. Activation of GPR40 potentiates GSIS [31,33,34] and increases secretion of the incretin hormones GLP-1, GIP and cholecystokinin [19,35-38] both ex vivo and in vivo. Recent studies suggest that in the β cell GPR40 primarily responds to endogenous fatty acids released in response to glucose stimulation and act in an autocrine manner to potentiate insulin secretion [39]. Although deletion of GPR40 is largely inconsequential for glucose metabolism under normal physiological conditions [33,40,41], high-fat diet (HFD) fed *gpr40*-null mice develop fasting hyperglycemia due to insufficient insulin secretion [42].

Complementarity between *gpr120* and *gpr40* tissue expression and function as well as overlapping ligands prompted us to investigate their possible interaction in the control of glucose homeostasis. Redundancy between *gpr120* and *gpr40* has been described in the gut where oral triglyceride-induced increases in plasma GIP and GLP-1 are dramatically reduced in double *gpr120/40* (Gpr120/40KO) but not *gpr120* (Gpr120KO) or *gpr40* (Gpr40KO) knockout mice [38]. Furthermore, coactivation of GPR120 and GPR40 in vivo with selective synthetic agonists or a dual agonist improves glucose control in *db/db* mice compared to activation of each receptor alone [43]. However, whether GPR120 and GPR40 cooperate to improve glucose control under physiological and pathophysiological conditions has not been addressed. In this study we investigated possible functional interactions between these two receptors in the control of glucose homeostasis with a particular emphasis on β-cell function. We generated single and double *gpr120* and *gpr40* KO mice and compared metabolic parameters, glucose and insulin tolerance as well as insulin secretion in vivo in both male and female mice fed a normal chow or HFD.

## MATERIALS AND METHODS

### Reagents and solutions

RPMI-1640 and FBS were from Life Technologies Inc. (Burlington, ON, Canada). Penicillin/Streptomycin was from Multicell Wisent Inc (Saint-Jean-Baptiste, QC, Canada). Fatty-acid-free BSA was from Equitech-Bio (Kerrville, TX, USA). Insulin (Humulin-R 100U/mL) was from Eli Lilly (Toronto, ON, Canada). Compound A (Cpd A) was from Cayman Chemical (Ann Arbor, MI, USA) and TAK-875 was from Selleckchem (Houston, TX, USA). All other reagents were from MilliporeSigma unless otherwise specified.

### Animals

All procedures involving animals were approved by the Institutional Committee for the Protection of Animals at the Centre Hospitalier de l’Université de Montréal. All mice were housed under controlled temperature on a 12h light/dark cycle with unrestricted access to water and laboratory chow. Mutant *gpr40* (RRID:MGI:3713765) [44] and *gpr120* (RRID:MGI:6477328) [45] colonies on a C57BL/6N (RRID:IMSR_JAX:005304) [46] background were maintained and genotyped as previously described. Male and female wild-type (WT, *gpr40*^+/+^;*gpr120*^+/+^), Gpr40KO (*gpr40*^-/-^;*gpr120*^+/+^), Gpr120KO (*gpr40*^+/+^;*gpr120*^-/-^) and Gpr120/40KO (*gpr40*^-/-^;*gpr120*^-/-^) experimental animals were generated in crosses between double heterozygotes (*gpr40*^+/-^;*gpr120*^+/-^). Animals were born at the expected Mendelian ratio. Both males and females were used for in vivo experiments and studied in separate cohorts. Animals were fed ad libitum with normal chow diet (CD; #2018 Teklad Global 18% protein rodent diet; 58% carbohydrate, 24% protein and 18% fat on a caloric basis; Harlan Teklad, Madison, WI) throughout the study or, after 8 weeks of age, switched to a HFD (#D12492; 60% fat, 20% protein and 20% carbohydrate on a caloric basis; Research Diets Inc., New Brunswick, NJ). Body weight, food intake and fed blood glucose were monitored weekly from 6-8 weeks of age.

### Metabolic tests

#### Oral Glucose Tolerance Tests (oGTT)

were performed on 4-6 hour fasted mice by measuring tail blood glucose and plasma insulin levels before (t=0 min) and 15, 30, 45, 60, 90 and 120 min after oral glucose administration (Dextrose 1g/kg BW). Tail blood glucose was measured using the hand-held glucometer Accu-Chek (Roche, Indianapolis, IN) and plasma insulin was measured by ELISA (Alpco Diagnostics; RRID:AB_2792981) [47] at 0, 15 and 30 min.

#### Intraperitoneal Insulin Tolerance Tests (ipITT)

Insulin (0.5 UI/kg, BW) was administered intraperitoneally on 4-6 hour fasted mice and tail blood glucose was measured before (t=0 min) and 15, 30, 45, 60, 90 and 120 min after insulin injection. Fasting insulinemia was measured by ELISA (Alpco Diagnostics; RRID:AB_2792981) [47] on plasma samples from tail vein blood collected before insulin injection.

#### Body weight composition

was assessed with the EchoMRI Analyzer-700 (Echo Medical System, Houston, TX).

#### Hyperglycemic clamps (HGC)

One-step HGC were performed in conscious, ad libitum fed animals as described [41]. Briefly, a 20% dextrose solution (Baxter, Mississauga, ON) was infused via a jugular catheter. Mice initially received a 90-second bolus (140 mg/kg/minute) and then the glucose infusion rate (GIR) was adjusted to maintain blood glucose between 14.5 and 17.7 mmol/l in females and between 15.7 and 19.7 mmol/l in males for 80 min. The insulin sensitivity index (M/I) was calculated as the glucose infusion rate (M) divided by the average plasma insulin during the last 30 min (i.e. from 50 to 80 min) of the clamp (I). The disposition index (DI) was calculated by multiplying the M/I by the average plasma C-peptide during the last 30 min of the clamp. Blood samples were collected from the tail to measure glucose, and plasma insulin (Alpco Diagnostics; RRID:AB_2792981) [47] and C-peptide (Alpco Diagnostics; RRID:AB_2801468) [48] were measured by ELISA at the time points indicated in the Figure legends.

### Islet isolation and static incubations for insulin secretion

Islets were isolated from 10-12 week-old male WT mice by collagenase digestion and dextran density gradient centrifugation as described previously [33] and recovered overnight in RPMI 1640 supplemented with 10% (wt/vol) FBS, 100 U/ml penicillin/streptomycin and 11 mM glucose. After recovery, triplicate batches of 20 islets each were incubated twice in KRBH (pH 7.4) with 0.1% (w/v) fatty-acid-free BSA and 2.8 mM glucose for 20 min., followed by a 1-hour static incubation in KRBH in the presence of 2.8 or 16.7 mM glucose. Insulin radioimmunoassay (MilliporeSigma; RRID:AB_2884035) [49] was used to measured insulin in the supernatant and intracellular insulin content after acid–alcohol extraction.

### Statistical analyses

Data are expressed as mean ± SEM. Significance was tested using ordinary one-way ANOVA or Brown-Forsythe and Welch ANOVA tests with their correction in cases of variance heterogeneity, two-way ANOVA with no assumption of sphericity and with post hoc adjustment for multiple comparisons or three-way ANOVA, as appropriate, using GraphPad Instat (RRID:SCR_000306) [50]. Tukey or Dunnett’s post hoc tests were performed as indicated in figure legends. P<0.05 was considered significant. Data points considered as outliers were excluded based on Grubbs’ test using the online calculator at: https://www.graphpad.com/quickcalcs/Grubbs1.cfm.

## RESULTS

### Lean Gpr120/40KO mice exhibit mild glucose intolerance

In a first cohort of animals, WT, Gpr40KO, Gpr120KO and double Gpr120/40KO male and female mice were CD-fed for 30 weeks. Metabolic parameters were assessed on a weekly basis and Echo-MRI, glucose and insulin tolerance tests, and HGC were performed as indicated (Fig. 1A). WT, Gpr40KO, Gpr120KO and double Gpr120/40KO male and female mice showed similar increases in body weight and energy intake over the course of the study (Fig. 1B-E). Percentages of fat and lean mass as assessed by Echo-MRI at 27-29 weeks of age were unaffected by genotype in male mice, whereas the proportion of fat to lean mass was slightly increased in female Gpr120KO compared to WT mice (Fig. 1F and G). Fasting blood glucose levels were significantly increased at 12 and 20 weeks of age in male and at all time points in female Gpr120/40KO compared to WT mice (Fig. 2A and B). A significant increase in fasting blood glucose compared to WT controls was also detected in male Gpr120KO mice at 20 weeks of age, in female Gpr40KO mice at 16, 20 and 24 weeks of age, and at all time points in female Gpr120KO mice. Weekly fed blood glucose levels were significantly increased in male Gpr40KO vs. WT mice and in female Gpr40KO and Gpr120/40KO mice vs. their WT and Gpr120KO littermates (Supplementary Fig. 1 in reference [51]).

**Figure 1.**
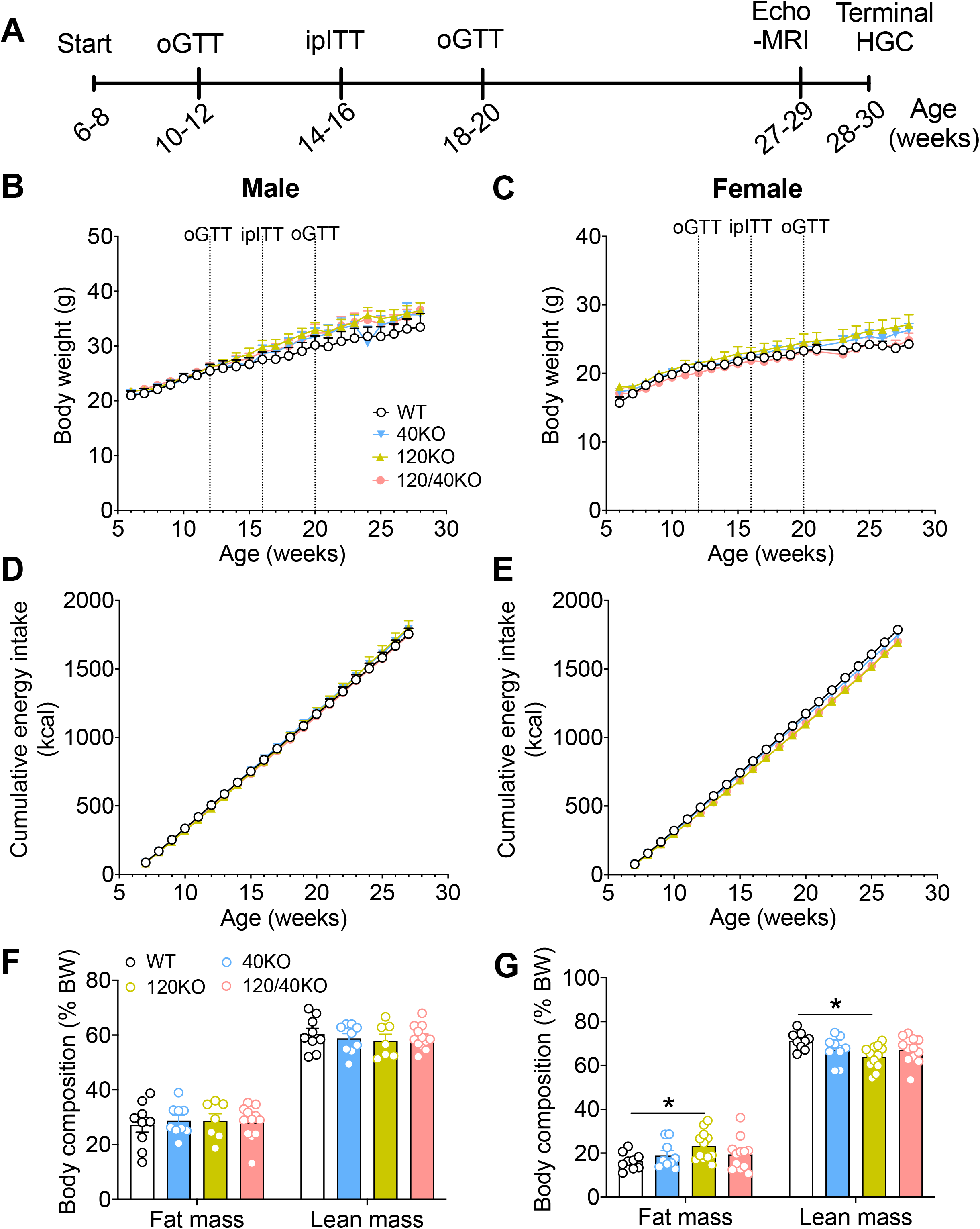
Energy balance in *gpr40* and *gpr120* single and double knockout (KO) mice under chow diet (CD). (**A**) Study plan to assess energy and glucose homeostasis in wild-type (WT), Gpr40KO (40KO), Gpr120KO (120KO) and Gpr120/40KO (120/40KO) male and female mice fed CD. (**B-G**) body weight (**B, C**), cumulative energy intake (**D, E**) and fat and lean mass as a percentage of body weight measured by Echo-MRI at 27-29 weeks of age (**F, G**) in male (**B, D, F**) and female (**C, E, G**) mice. Data are presented as mean +/- SEM (n=6-8 animals per group). P-values were calculated by regular 2-way ANOVA or mixed model when some follow-up values were missing with no assumption of sphericity (**B-E**) or 2-way ANOVA with post-hoc Tukey’s multiple comparisons test (**F, G**). *p < 0.05.

**Figure 2.**
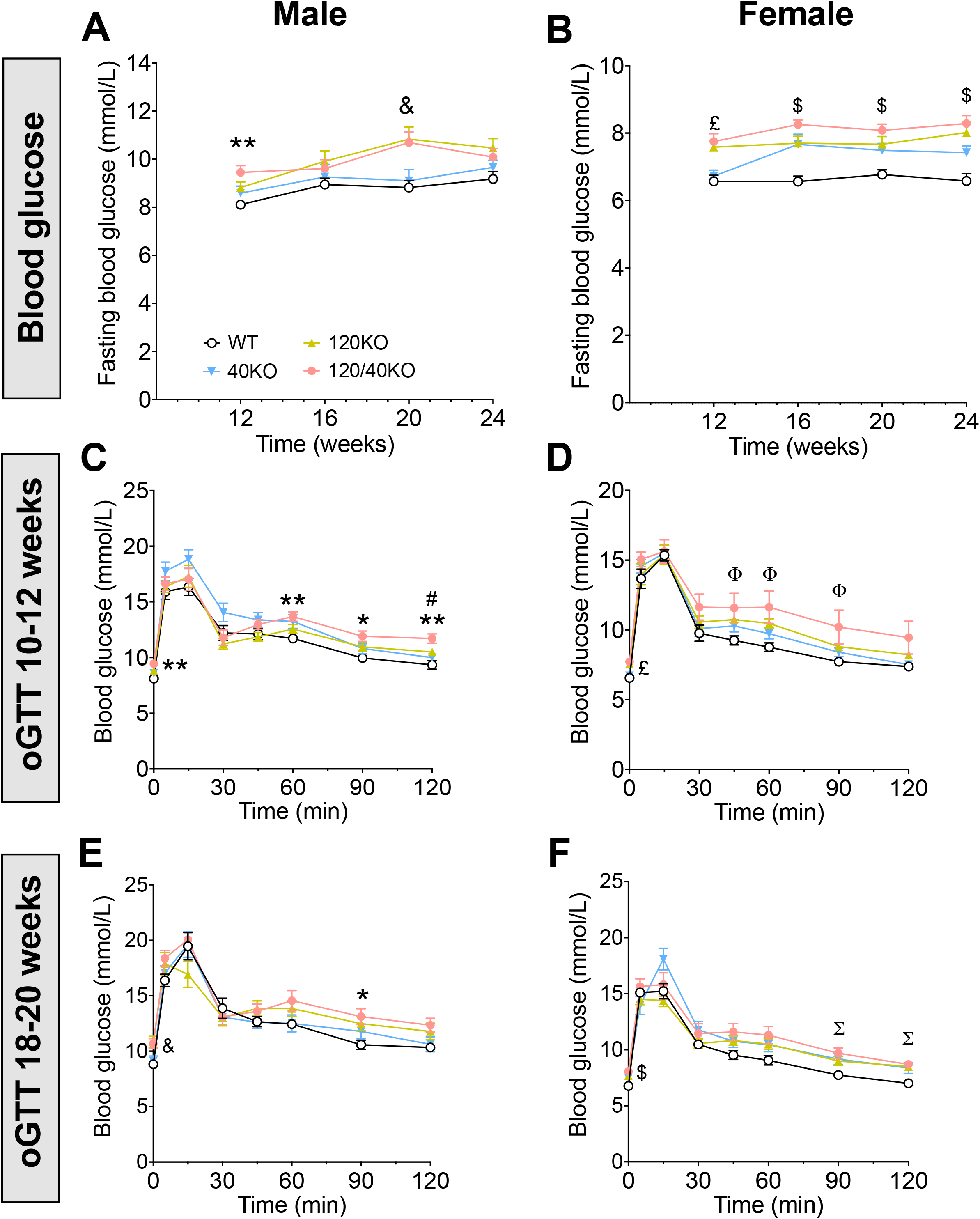
Glucose homeostasis in *gpr40* and *gpr120* single and double knockout (KO) mice under chow diet (CD). (**A, B**) Fasted blood glucose at 12, 16, 20 and 24 weeks of age in wild-type (WT), Gpr40KO (40KO), Gpr120KO (120KO) and Gpr120/40KO (120/40KO) male (**A**) and female (**B**) mice fed CD. (**C-F**) Blood glucose during an oGTT (glucose 1g/kg BW at 0 min) at 10-12 (**C, D**) and at 18-20 (**E, F**) weeks of age in male (**C, E**) and female (**D, F**) mice. Data are presented as mean +/- SEM (n=7-13 animals per group). P-values were calculated by regular 2-way ANOVA or mixed model when some values were missing, with no assumption of sphericity and with Tukey’s multiple comparisons tests. **p<0.01 and *p<0.05, 120/40KO vs WT; &, p<0.05 120/40KO and 120KO vs WT; £, p<0.01 120/40KO and 120KO vs WT, p<0.05 120/40KO and 120KO vs Gpr40KO; $, at least p<0.05 for 120KO and 40KO vs WT and at least p<0.005 for the 120/40KO vs WT; #, p<0.05 120/40KO vs 40KO; Φ, at least p<0.05 WT vs 120KO; Σ, at least p<0.05 120/40KO and 120KO vs WT.

oGTT performed at 10-12 weeks of age revealed prolonged glucose excursions in male Gpr120/40KO mice that were significantly different between 60-120 min post oral glucose load compared to WT controls and at 120 min compared to Gpr40KO mice (Fig. 2C). In females, both single and double Gpr120/40KO mice exhibited reduced glucose tolerance between 45-90 min post oral glucose administration compared to WT mice, but the difference was significant only for the Gpr120KO mice (Fig. 2D). oGTT performed at 18-20 weeks of age revealed mild glucose intolerance in male Gpr120/40KO mice at 90 min and in female Gpr120/40KO and Gpr120KO mice between 90-120 min compared to WT controls (Fig. 2E and F). Plasma insulin levels determined during the oGTT at 10-12 and 18-20 weeks of age revealed no significant differences between genotypes in either male or female mice (Supplementary Fig. 2 in reference [51]).

ipITT performed in male and female mice at 14-16 week of age revealed no significant difference in insulin tolerance between genotypes (Supplementary Fig. 2 in reference [51]).

We performed HGC at 28-30 weeks of age to assess β cell function in vivo. Blood glucose levels were within the target range between 50 and 80 min of the clamp (Fig. 3A and B). Although plasma insulin (Fig. 3C and D) and C-peptide (Fig. 3E and F) levels were similar in the 4 genotypes, the GIR and DI were significantly lower in male Gpr120/40KO vs. Gpr40KO mice (Fig. 3G and I). A similar, albeit not statistically significant trend was observed in female mice (Fig. 3H and J). Given that the reduction in DI is mostly driven by a lower GIR without significant changes in c-peptide levels, we conclude from these data that the combined deletion of *gpr120* and *gpr40* does not significantly impair insulin secretion in vivo. The absence of significant differences between Gpr120/40KO and Gpr120KO mice suggests that deletion of *gpr120* accounts for the majority of the observed phenotype.

**Figure 3.**
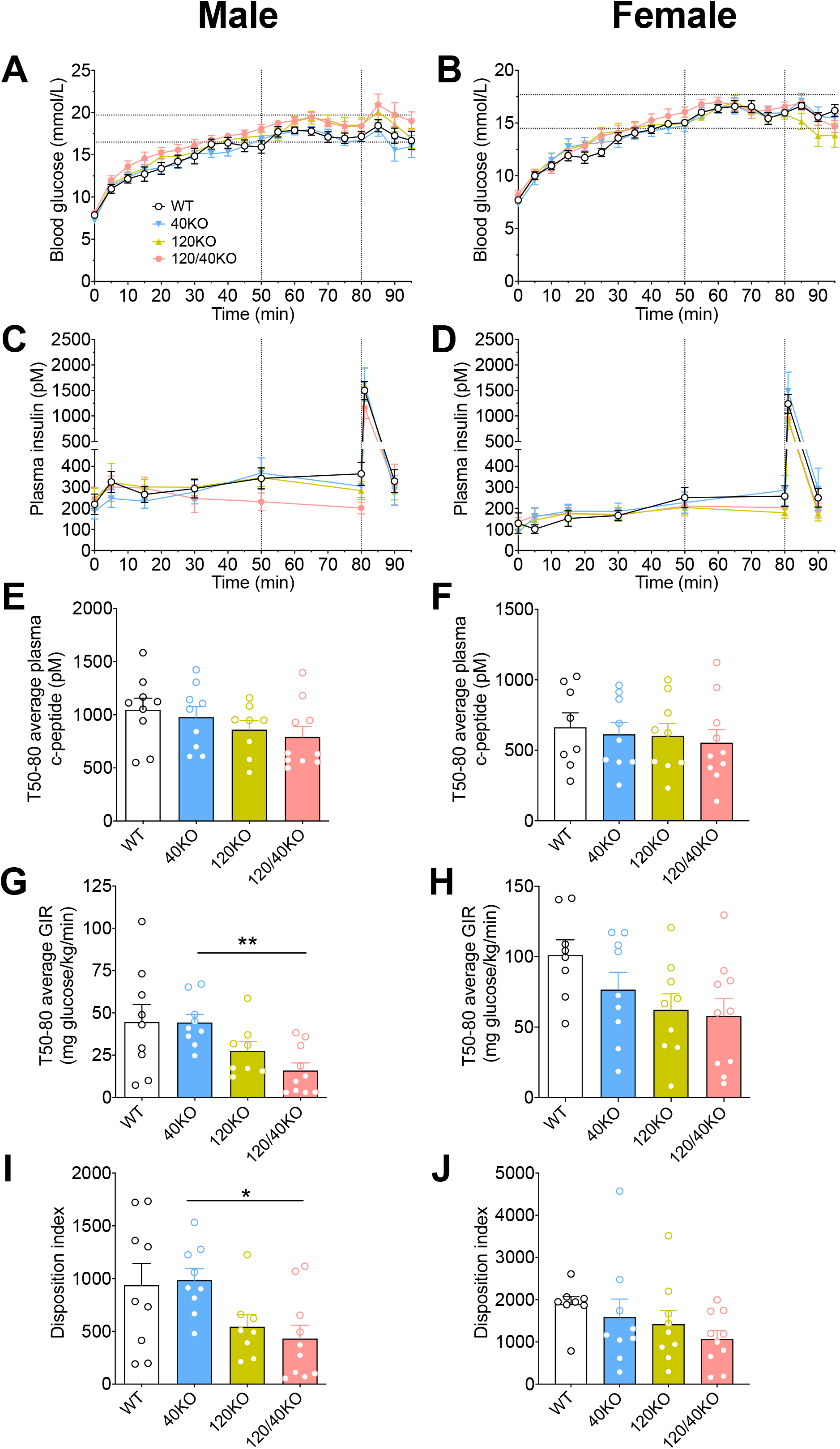
Hyperglycemic clamp analysis of *gpr40* and *gpr120* single and double knockout mice under chow diet. (**A-D**) Blood glucose (**A, B**) and plasma insulin (**C, D**) during the hyperglycemic clamp (HGC) in male (**A, C**) and female mice (**B, D**). (**E-J**) Plasma c-peptide (**E, F**) and glucose infusion rate (GIR) (**G, H**) during the steady state (50-80 min) and disposition Index (DI) (**I, J**) in male (**E, G, I**) and female (**F, H, J**) mice. Data are presented as mean +/- SEM (n=8-10 animals per group). (**A-D**) P-values were calculated by the mixed model of 2-way ANOVA with Greisser-Greenhouse correction in case of variance heterogeneity (i.e. no assumption of sphericity) and with Tukey’s post-hoc multiple comparisons tests. No significant effect of genotype on blood glucose or plasma insulin levels was detected during the steady state (50-80 min) in both males and females. (**E-J**) P-values were calculated by Brown-Forsythe and Welch’s ANOVA (**G**) or ordinary one-way ANOVA (**E, F, H-J**) with Dunnett’s T3 or Tukey’s post-hoc MCTs, respectively. **p<0.01, *p<0.05.

### High-fat fed Gpr120/40KO mice have normal glucose tolerance

In a second cohort of animals, WT, Gpr40KO, Gpr120KO and Gpr120/40KO male and female mice were fed either CD or HFD beginning at 8 weeks of age for a total of 12 weeks. Metabolic parameters were assessed on a weekly basis. Echo-MRI and oral glucose tolerance tests were performed as indicated (Fig. 4A). Both male and female HFD-fed mice accumulated more weight than CD-fed animals, as expected, but no differences were detected between genotypes on either feeding regimen (Fig. 4B and C). Male Gpr120/40KO mice fed a HFD exhibited reduced energy intake compared to Gpr120KO and WT controls (Fig. 4D). In contrast, females showed no differences in energy intake between genotype following either CD or HFD (Fig. 4E). Relative fat mass was increased and lean mass decreased in HFD fed compared to CD-fed male and female animals but no differences were detected between genotypes (Fig. 4F and G).

**Figure 4.**
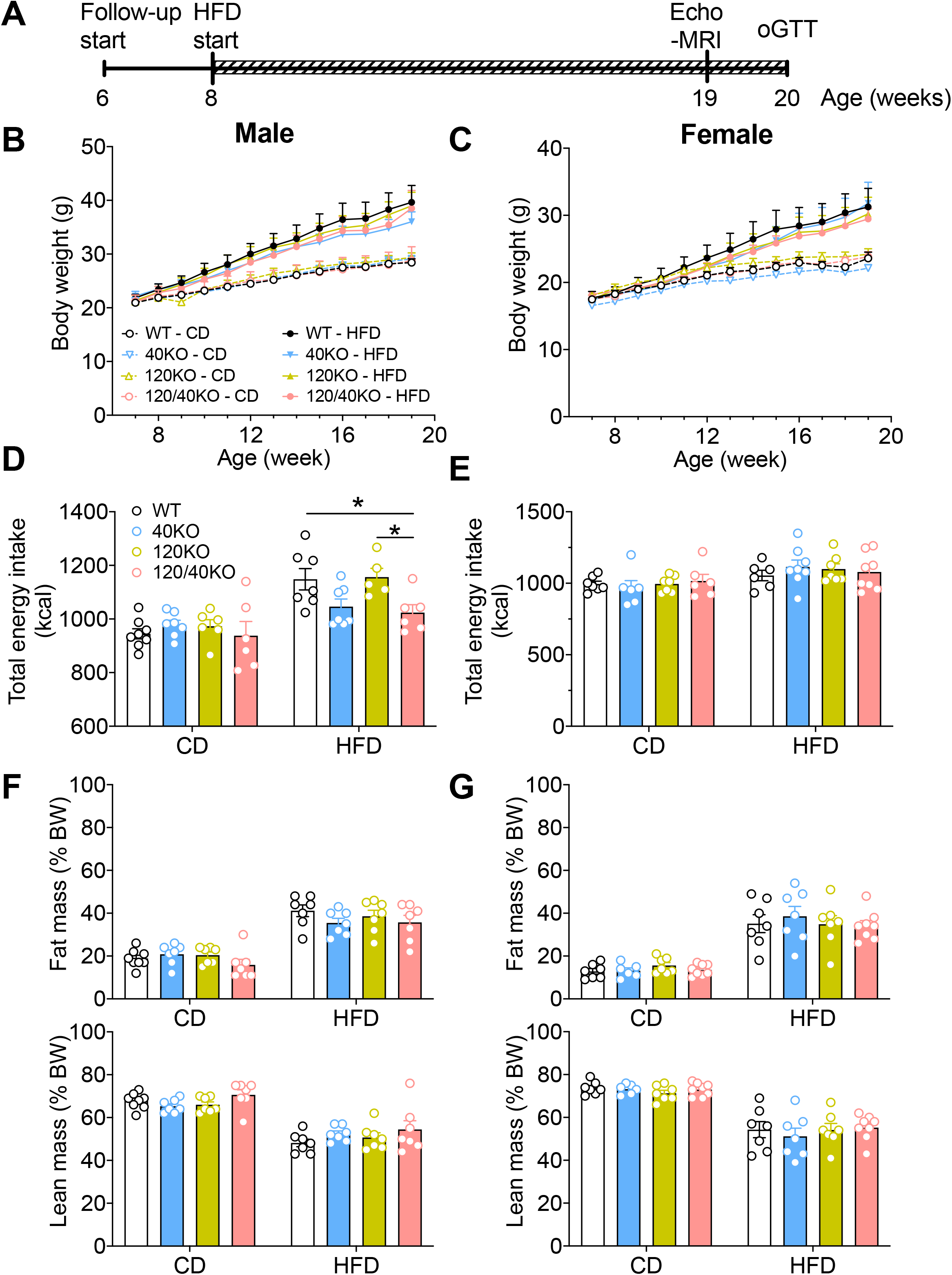
Energy balance in *gpr40* and *gpr120* single and double knockout (KO) mice under chow (CD) or high-fat diet (HFD). (**A**) Study plan to assess energy and glucose homeostasis in wild-type (WT), Gpr40KO (40KO), Gpr120KO (120KO) and Gpr120/40KO (120/40KO) male and female mice fed CD or HFD from 8 to 20 weeks of age. (**B-G**) body weight (**B, C**), total energy intake at 20 weeks of age (**D, E**) and fat and lean mass as a percentage of body weight measured by Echo-MRI at 19 weeks of age (**F, G**) in male (**B, D, F**) and female (**C, E, G**) mice. Data are presented as mean +/- SEM (n=7-10 animals per group). P-values were calculated by regular 2-way ANOVA or mixed model when some follow-up values were missing with no assumption of sphericity (**B, C**) or 2-way ANOVA with post-hoc Tukey’s multiple comparisons test (**D-G**). *p < 0.05.

Mean and/or weekly fed blood glucose levels throughout the study were increased in CD-fed male Gpr120/40KO compared to all other genotypes and in HFD-fed Gpr120/40KO and Gpr120KO compared to WT mice but unaffected by genotype in female mice (Supplementary Fig. 3 in reference [51]).

oGTT performed at 20 weeks of age revealed reduced glucose tolerance in male Gpr120/40KO mice fed a CD at 60 and 120 min compared to WT controls (Fig. 5A), similar to our previous findings (Fig. 2). However, plasma insulin levels determined during the test revealed no significant differences between genotypes (Fig. 5B). A non-significant trend for reduced glucose tolerance was observed in Gpr120/40KO females (Fig. 5C), without changes in plasma insulin levels (Fig. 5D). Following HFD, no significant differences in glucose tolerance or plasma insulin levels were detected between genotypes in either male (Fig. 5E and F) or female (Fig. 5G and H) mice. Taken together these data suggest that combined deletion of *gpr120* and *gpr40* does not aggravate glucose intolerance under HFD in either male or female mice.

**Figure 5.**
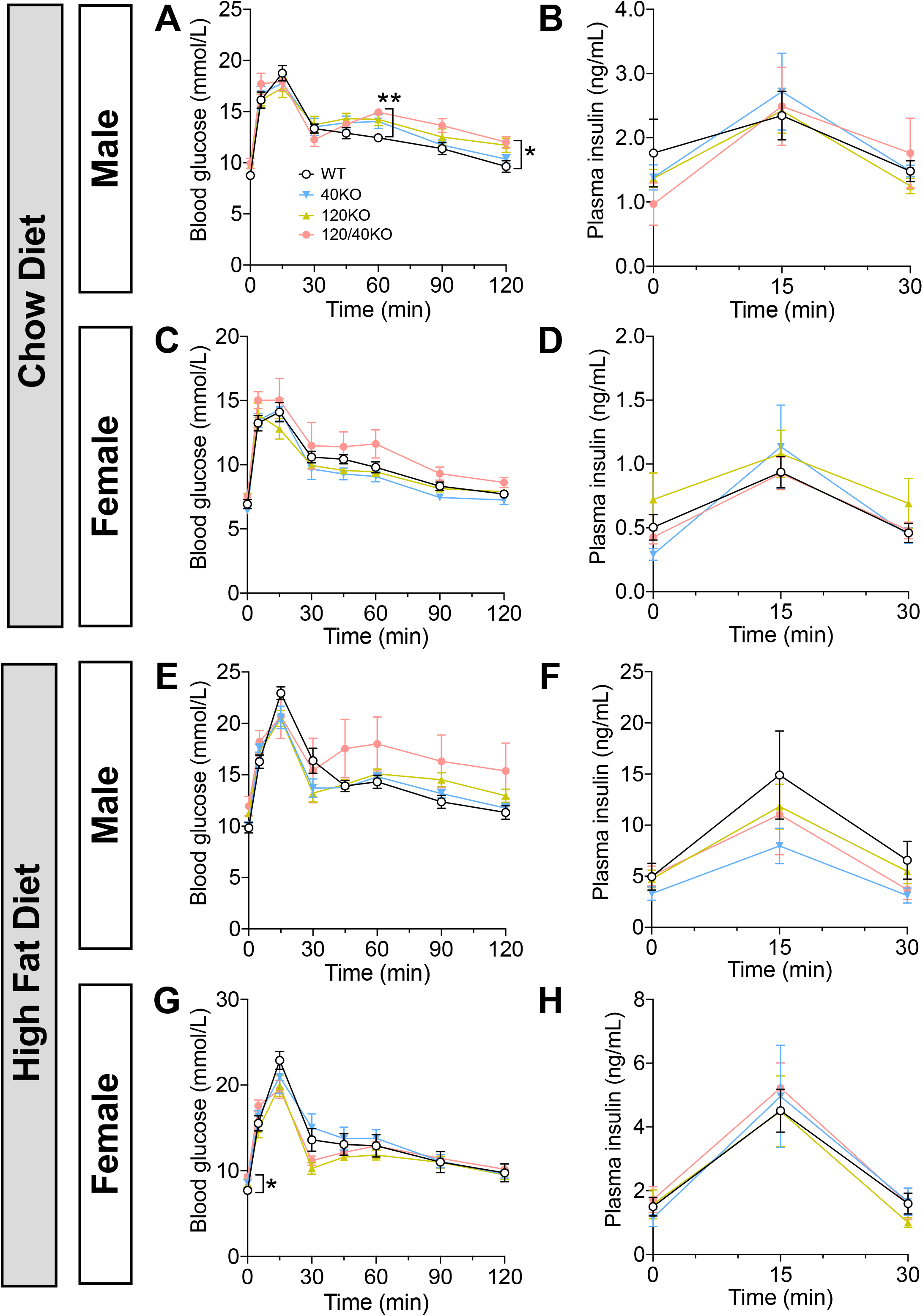
Oral glucose tolerance in *gpr40* and *gpr120* single and double knockout (KO) mice under chow (CD) or high-fat diet (HFD). (**A-H**) Blood glucose (**A, C, E, G**) and plasma insulin (**B, D, F, H**) levels during an oGTT (glucose 1g/kg BW at 0 min) at 20 weeks of age in male (**A, B, E, F**) and female (**C, D, G, H**) wild-type (WT), Gpr40KO (40KO), Gpr120KO (120KO) and Gpr120/40KO (120/40KO) mice fed CD (**A-D**) or HFD (**E-H**) from 8 to 20 weeks of age. Data are presented as mean +/- SEM (n=6-8 animals per group). P-values were calculated by regular 2-way ANOVA or mixed model when some values were missing, with no assumption of sphericity and with Tukey’s multiple comparisons tests. **p<0.01 and *p<0.05, 120/40KO vs WT.

### GPR120 and GPR40 agonists additively potentiate GSIS in isolated mouse islets

Agonists of GPR120 or GPR40 enhance GSIS in isolated mouse islets [23-26,30,31,33,34]. Here we asked whether simultaneous activation of GPR120 and GPR40 potentiates GSIS additively or synergistically compared to the activation of each receptor alone. Islets were isolated from lean male WT mice and insulin secretion was assessed in response to the selective GPR40 agonist TAK-875 and GPR120 agonist Cpd A, singly or in combination (Fig. 6). Glucose alone increased insulin secretion (Fig. 6A and B), as expected. TAK-875 and Cpd A alone weakly but significantly potentiated this effect and combined exposure to TAK-875 and Cpd A additively increased GSIS (Fig. 6A and B). No significant differences were detected in total insulin content (Fig. 6C).

**Figure 6.**
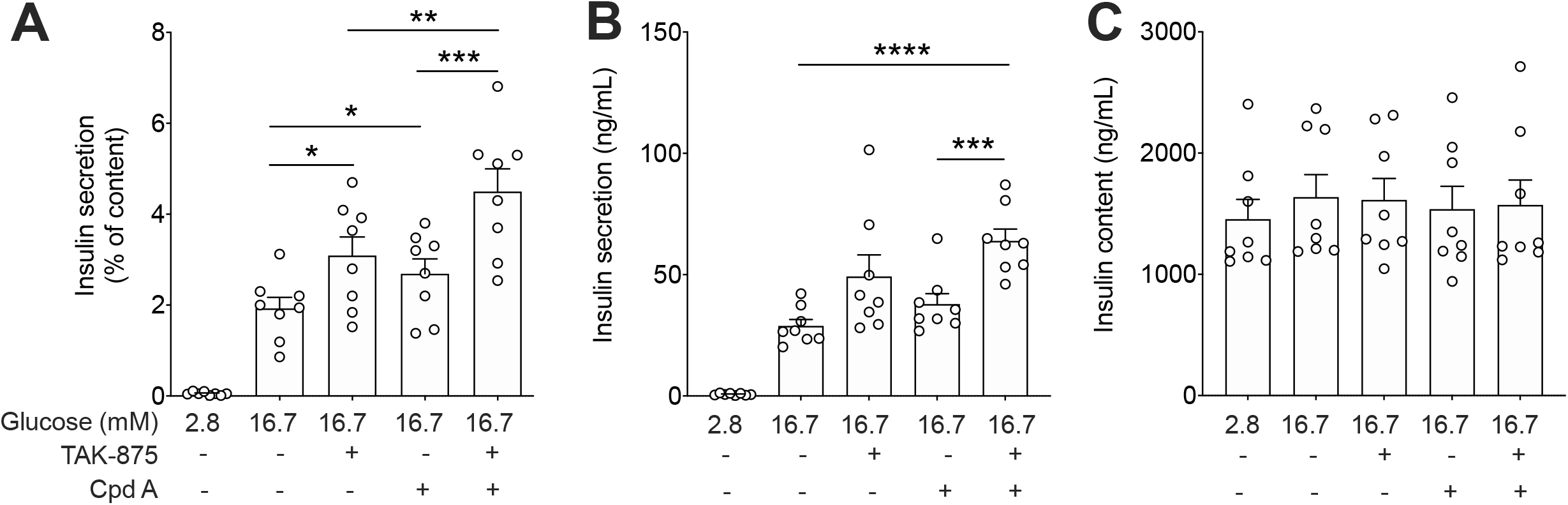
Potentiation of GSIS in response to Cpd A and TAK-875 in islets ex vivo. Insulin secretion was assessed in 1-h static incubations in response to 2.8 or 16.7 mM glucose with or without the GPR120 agonists Cpd A (50 µM) and the GPR40 agonist TAK-875 (5 µM). **(A-C)** Insulin secretion presented as a percentage of islet insulin content (**A**), insulin secretion (**B**) and insulin content (**C**). Data represent individual values and are expressed as mean +/- SEM of 6 independent experiments. P-values were calculated by regular two-way ANOVA with Dunnett’s or Tukey’s (G-J) post hoc adjustment for multiple comparisons and Welch/Brown-Forsythe correction when necessary to compensate for SD variances. * p<0.05, ** p<0.005, *** p<0.001, **** p<0.0001.

## DISCUSSION

The objective of this study was to assess possible interactions between GPR120 and GPR40 in the control of glucose homeostasis and β cell function under physiological conditions and metabolic stress. By comparing adult, chow-fed WT and *gpr120* and *gpr40* single and double KO mice we showed that the combined loss of both genes leads to a slight reduction in glucose tolerance but no substantial impairment of insulin secretion in vivo. In most cases, the Gpr120/40KO mice phenotype seemed to be mainly related to the loss of GPR120 and slightly aggravated by the loss of GPR40, suggesting that the two receptors signaling pathways act additively rather than synergistically. No significant differences between genotypes were detected in glucose homeostasis following 12 weeks of high-fat feeding in either sex. We then showed that selective agonists for GPR120 and GPR40 additively potentiate GSIS in isolated islets ex vivo.

### Gpr120KO

Deletion of *gpr120* had no significant impact on body weight in CD-fed male or female mice, in agreement with previous studies [3,5]. In contrast, Suckow et al. [27] observed an increase in body mass in *gpr120*-null mice following a similar diet regimen. In our study, fat mass was increased in female Gpr120KO mice which may be accounted for by the reduced metabolic rate, inhibition of adipocyte lipolysis and increased size of adipocytes observed previously in *gpr120*-null mice [5,43].

Fasting glucose levels were elevated in male and female Gpr120KO mice under CD, but oral glucose tolerance was unaffected in male and mildly reduced in female Gpr120KO mice, and there were no differences in circulating insulin levels or insulin tolerance. These findings agree with previous reports [5,24]. They do, however, contrast with some studies which found glucose intolerance [3,27] and elevated plasma insulin levels and insulin resistance [3] in *gpr120*-null male mice. Age-related differences may play a role as insulin tolerance test were performed at 14-16 weeks of age in this study whereas Oh et al., analyzed mice using hyperinsulinemic/euglycemic clamps at 23-28 weeks of age [3].

Following HFD, body weight, glucose tolerance and insulin levels were not affected in either male or female Gpr120KO mice, similar to previous findings in male mice [3,4,52]. This is in contrast to the findings of Ichimura et al. [5] who reported that HFD-fed *gpr120*-null male mice gain more weight and are glucose intolerant. Discrepancies in *gpr120*-null metabolic phenotypes may be accounted for by differences in diet composition and genetic background, such as the mixed C57Bl/6/129 used by Ichimura et al. [5] vs. the pure C57Bl/6N used by Bjursell et al. [52] and in this study. It should be noted, however, that a role of GPR120 signaling has been demonstrated in the context of chronic administration of specific agonists such as Cpd A [4,6,26] and GSK137647 [26] which attenuate the negative effects of HFD on energy and glucose homeostasis in WT but not *gpr120*-null mice. This suggests that during metabolic stress and/or chronic feeding with a diet poor in omega-3 fatty acids, levels of endogenous GPR120 ligands may be insufficient to confer a protective effect.

### Gpr40KO

Deletion of *gpr40* alone had no significant impact on energy homeostasis in male or female mice fed either CD or HFD, as shown previously [33,41,42,53].

Glucose and insulin tolerance and insulin secretion in CD-fed Gpr40KO mice were also largely unaffected, as previously observed [33]. Contrary to our previous findings [42], we did not detect significant differences in fasting glucose levels in chow-fed, male Gpr40KO mice vs WT. This could be accounted for by a greater statistical power in our previous study, in which only 2 groups with a larger sample size were compared, and/or differences in the age of the mice (7 weeks in [42] vs 12-24 weeks in this study). In the present study, we found however that female Gpr40KO mice had higher fasting glucose, and male and female mice had higher fed glucose, than WT controls. The significant reduction in insulin levels previously detected during HGC in *gpr40*-null mice [41] was not seen in the present study, which may be due to difference in the age and/or strain of the animals. The nicotinamide nucleotide transhydrogenase mutation and other polymorphisms in the C57Bl/6J background used in our previous study [41] – vs. the C57Bl/6N background used here - are known to attenuate insulin secretion [54] and may interact with the *gpr40* mutation to aggravate the secretory defect.

In a HFD study by Kebede et al. [42] we observed a precocious increase in fasting blood glucose levels after 3 weeks of diet in male Gpr40KO mice which normalized after 8 weeks, similar to our findings after 12 weeks of diet.

### Gpr120/40KO

Although weight gain was similar, male, but not female, Gpr120/40KO mice exhibited reduced energy intake compared to Gpr120KO and WT controls. GPR120 and GPR40 are expressed in hypothalamic microglia and neurons, respectively [55]. Interestingly, intracerebroventricular injection of GW9508, a GPR120 and GPR40 dual agonist, increases food intake in HFD fed mice [55]. Although the underlying reason for the decreased food intake in Gpr120/40KO mice is unknown, loss-of-function of these genes in the hypothalamus would be expected to alter food intake.

In mice fed a CD, we observed an increase in fasting glucose levels in both male and female Gpr120/40KO and a slight decrease in glucose tolerance in males. These changes were not observed in any of the groups under HFD. The reduction in glucose tolerance was more pronounced in double Gpr120/40KO compared to single Gpr120KO or Gpr40KO mice, indicative of an additive effect. As no differences were detected in insulin tolerance between genotypes, the mild reduction in glucose tolerance in Gpr120/40KO mice might be due to deletion of the genes in pancreatic islets and/or reduced incretin hormone secretion. Along these lines, Ekberg et al. [38] demonstrated that *gpr120* and *gpr40* are highly expressed in GLP-1 and GIP-secreting cells in the gut, and that the combined loss of the two receptors leads to a major reduction in GLP-1 and GIP levels following oral triglyceride administration, which is not seen in mice deficient in each receptor alone. However, we cannot exclude that a mild insulin resistance in Gpr120KO and Gpr120/40KO compared to WT mice, not detected in the ipITT, contributes to the increased fasting glucose levels and glucose intolerance. This possibility is consistent with the lower GIR observed during HGC in 28-30 weeks Gpr120/40KO mice, and is supported by the finding that Gpr120KO mice can develop insulin resistance [3]. However, glucagon secretion and glucagon sensitivity are elevated in Gpr120KO mice [27] and Gpr40 agonist treatment inhibits glucose production in Goto Kakizaki rat through changes in gluconeogenesis [56]. Hence, increased glycogenolysis and gluconeogenesis could also contribute to higher blood glucose levels in Gpr120/40KO mice.

The reduced DI in HGC was largely due to a reduction in GIR with no changes in insulin or c-peptide levels in male Gpr120/40KO vs single Gpr40KO. Nevertheless, the reduced GIR is suggestive of a relative functional β-cell deficit, possibly due to a lack of β-cell mass compensation to age-related insulin resistance in Gpr120/40KO males. Indeed, *gpr120* deletion is associated with insulin resistance in mice of a similar age [3]. Although our previous studies indicate that *gpr40* deletion alone does not impact β-cell mass [41], given the role of GPR120 and GPR40 in the regulation of hormones that control β-cell proliferation, including somatostatin [20,28,30] and GLP-1 [16,24,35,36,38], it is possible that their deletion impacts β-cell mass. We did not test for this possibility in our study. The reduced GIR seems to be mainly attributable to the deletion of *gpr120* and slightly aggravated by the additional loss of *gpr40*. A non-significant trend was detected in female Gpr120/40KO mice. Phenotypic differences between male and female mice may be due to a number of sex-related factors known to contribute to sex differences in metabolism [57].

Alterations in glucose homeostasis in Gpr120/40KO mice following CD but not HFD are notable for their similarities to Gpr120KO mice, which show the same trends. We surmise that HFD may diminish GPR120 signaling thereby limiting the effect of gene loss in single and double Gpr120KO mice compared to Gpr40KO and WT animals. This could be in part related to lower intake of GPR120 ligands, since HFD are poor in ω-3 polyunsaturated FAs compared to CD.

### Isolated islets

The stimulatory effects of the GPR120 agonist Cpd A and the GPR40 agonist TAK-875 were additive and not synergistic. This is consistent with our in vivo observations in double KO mice and suggests that the signaling mechanisms of the two receptors in islets are independent. Recently, using a δ-cell specific Gpr120KO mouse we showed that Cpd A potentiates GSIS largely via δ cell-specific GPR120 inhibitory signaling that dampens cAMP generation and somatostatin secretion, alleviating a paracrine inhibitory signal that restricts insulin secretion [30]. In contrast, GPR40 is predominantly expressed in islet β cells where its activation directly promotes insulin secretion via Gαq/11 coupling and phospholipase C – protein kinase D1 signaling [34]. However, our studies also revealed that *gpr120* is expressed in β cells and that Cpd A increases cAMP generation and calcium fluxes in β cells [30]. Furthermore, in a recent study, ciliary GPR120 activation was shown to increase cAMP levels and promote insulin secretion in intact islets and insulin-secreting cell lines [58]. Together these studies suggest that although GPR120 inhibitory signaling in δ cells indirectly increases insulin secretion, GPR120 stimulatory signals in β cells may also contribute to its modulation of islet hormone secretion. This model is consistent with our observation that the stimulatory effects of GPR120 and GPR40 on insulin secretion are additive rather than synergistic.

## Conclusion

Our results show that combined deletion of GPR120 and GPR40 minimally alters glucose homeostasis in mice, with only a mild impairment of glucose tolerance in male animals. Our data also suggest that the two receptors signaling cooperate in an additive rather than in a synergistic manner to potentiate glucose-stimulated insulin secretion ex vivo. These findings contribute to our understanding of the role of these fatty-acid GPCR in the regulation of glucose homeostasis and islet function.

## ACKNOWLEDGEMENTS

We thank Annie Levert (CRCHUM) for valuable technical assistance.

## DATA AVAILABILITY

The datasets generated during and/or analyzed during the current study are not publicly available but are available from the corresponding author on reasonable request.

## Notes

https://doi.org/10.6084/m9.figshare.13365962.v1

### Competing Interest Statement

The authors have declared no competing interest.

