## Supplementary Figures for "Combined deletion of free fatty-acid receptors 1 and 4 minimally impacts glucose homeostasis in mice"

### Supplementary Figure 1

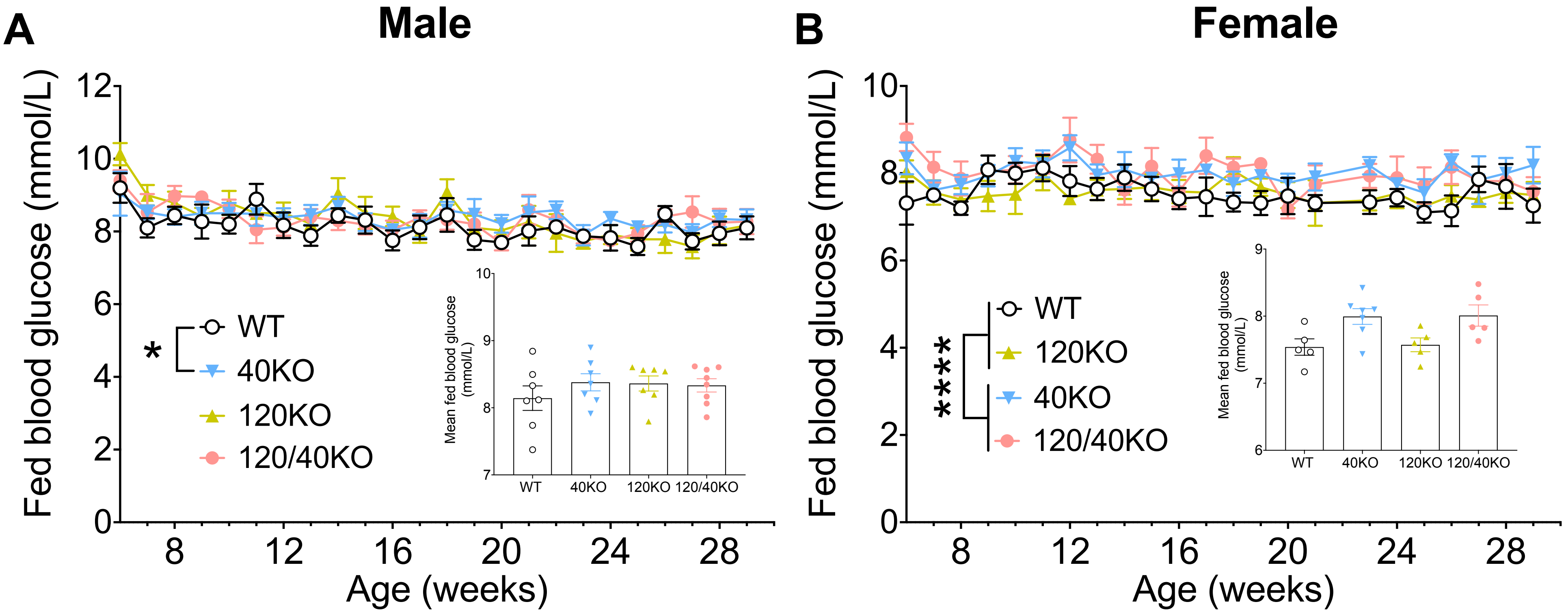


**Supplementary Figure 1. Fed blood glucose levels in Gpr40 and Gpr120 single and double knockout (KO) mice under chow diet (CD).** (**A, B**) weekly and mean (insert) fed blood glucose throughout the study in wild-type (WT), Gpr40KO (40KO), Gpr120KO (120KO) and Gpr120/40KO (120/40KO) male (**A**) and female (**B**) mice fed CD. Data expressed as mean +/- SEM of 5-8 independent experiments. P-values were calculated between groups over the entire study period by regular 2-way ANOVA or mixed model when some values were missing, with no assumption of sphericity for weekly fed blood glucose and by One-way ANOVA for mean fed blood glucose (insert), both with Tukey's multiple comparisons tests. *p<0.05, ****p<0.0001.

### Supplementary Figure 2

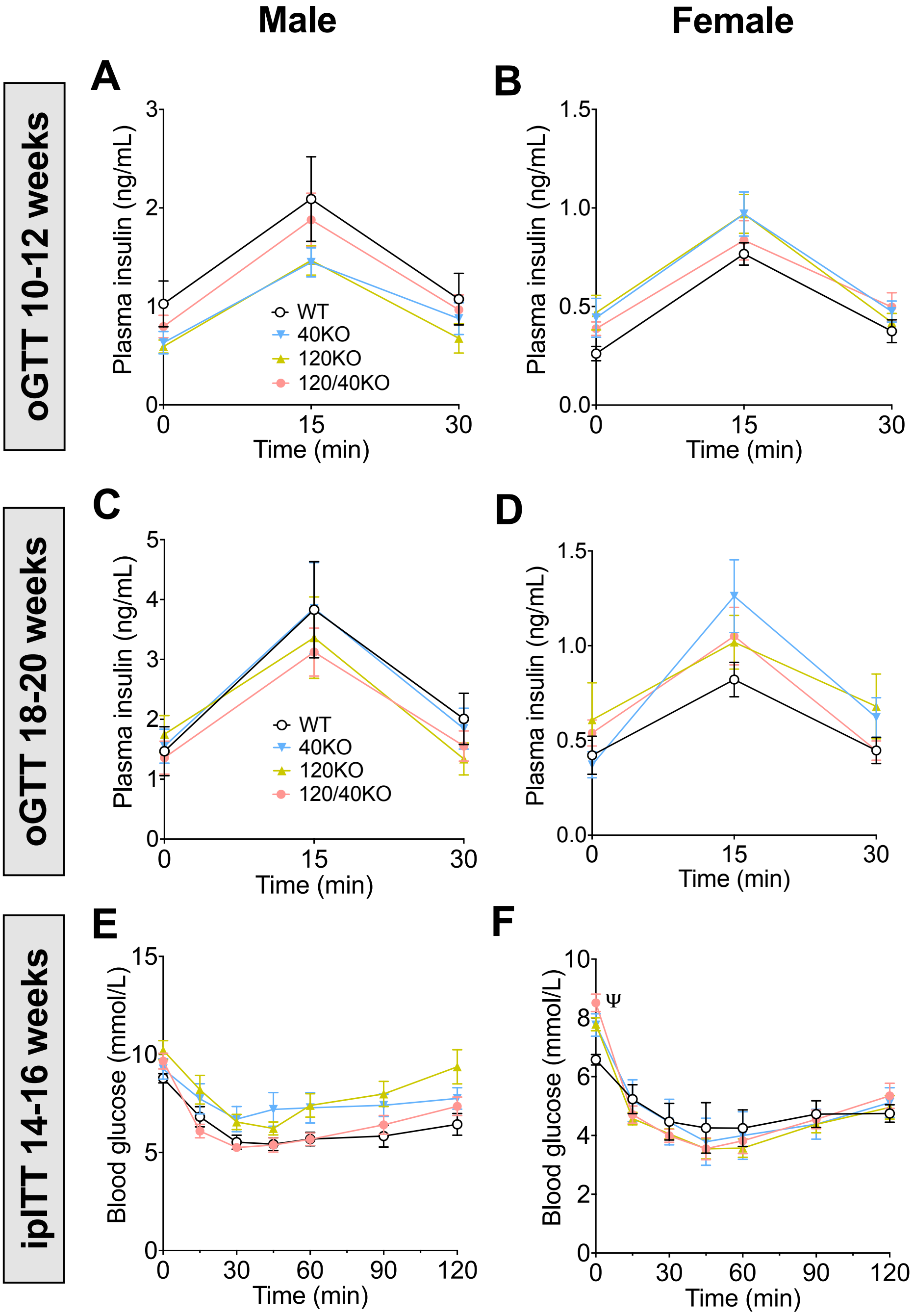


**Supplementary Figure 2. Plasma insulin levels during oGTT and blood glucose levels during ipITT in Gpr40 and Gpr120 single and double knockout (KO) mice under chow diet (CD).** (**A-F**) Plasma insulin levels during an oGTT (glucose 1g/kg BW at 0 min) at 10-12 (**A, B**) and 18-20 (**C, D**) weeks of age and blood glucose during an ipITT (insulin 0.5 UI/kg BW at 0 min) at 14-16 weeks of age (**E, F**) in wild-type (WT), Gpr40KO (40KO), Gpr120KO (120KO) and Gpr120/40KO (120/40KO) male (**A, C, E**) and female (**B, D, F**) mice fed CD. Data are presented as mean +/- SEM (n=7-13 animals per group). P-values were calculated by regular 2-way ANOVA or mixed model when some values were missing, with no assumption of sphericity and with Tukey's multiple comparisons tests. ψ, $p<0.01$ 120KO vs WT and $p<0.005$ 120/40KO vs WT.

### Supplementary Figure 3

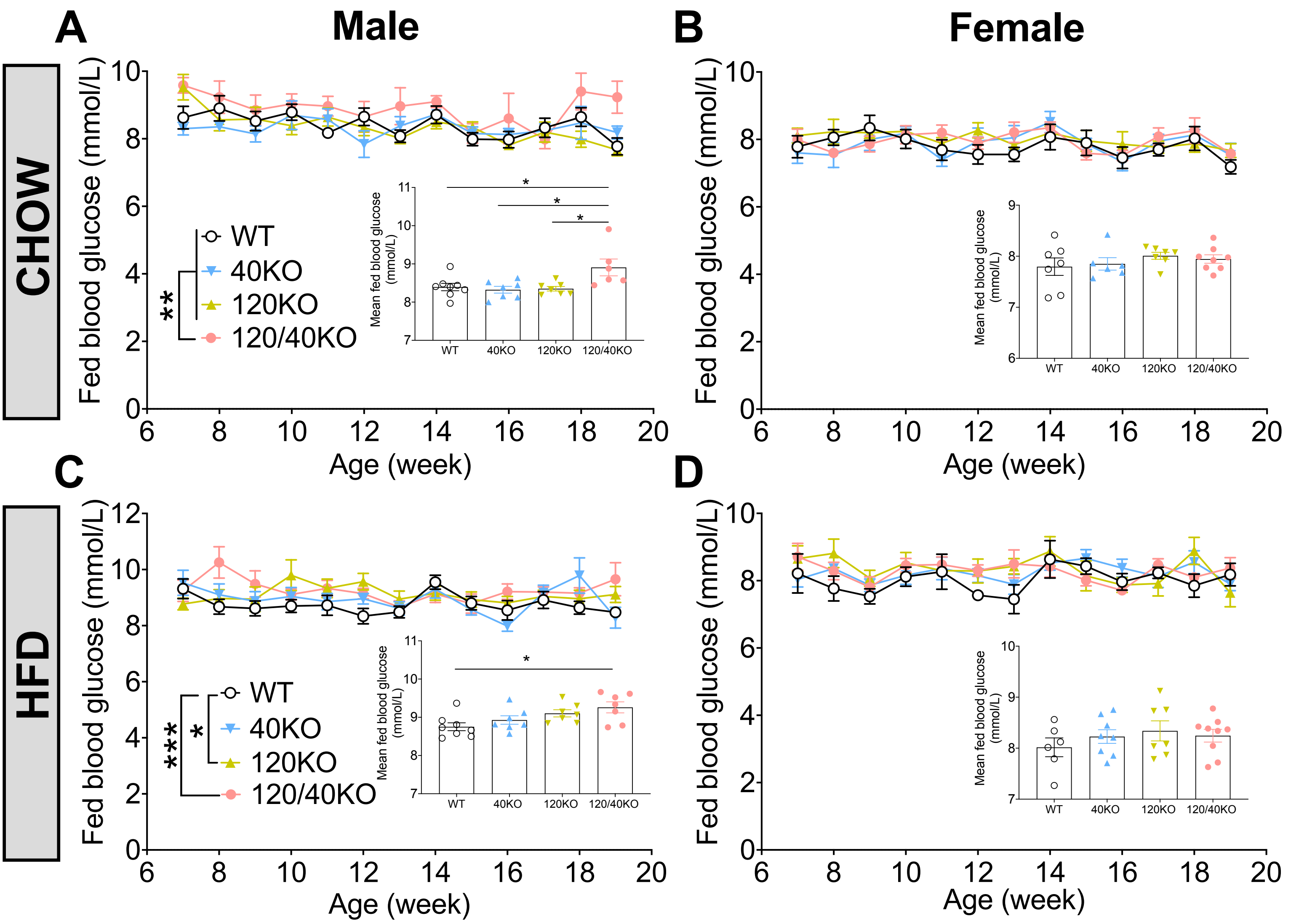


**Supplementary Figure 3. Fed blood glucose levels in Gpr40 and Gpr120 single and double knockout (KO) mice under chow (CD) or high-fat diet (HFD).** (**A-D**) Weekly and mean (insert) fed blood glucose throughout the study in wild-type (WT), Gpr40KO (40KO), Gpr120KO (120KO) and Gpr120/40KO (120/40KO) male (**A, C**) and female (**B, D**) mice fed CD (**A, B**) or HFD (**C, D**). Data expressed as mean +/- SEM of 6-9 independent experiments. P-values were calculated between groups over the entire study period by regular 2-way ANOVA or mixed model when some values were missing, with no assumption of sphericity for weekly fed blood glucose and by One-way ANOVA for mean fed blood glucose (insert), both with Tukey's multiple comparisons tests : *p<0.05, **p<0.01, ***p<0.001.
